# Multi-attribute Glycan Identification and FDR Control for Glycoproteomics

**DOI:** 10.1101/2021.10.29.466473

**Authors:** Daniel A. Polasky, Daniel J. Geiszler, Fengchao Yu, Alexey I. Nesvizhskii

**Affiliations:** Department of Pathology, University of Michigan; Department of Computational Medicine and Bioinformatics, University of Michigan

## Abstract

Rapidly improving methods for glycoproteomics have enabled increasingly large-scale analyses of complex glycopeptide samples, but annotating the resulting mass spectrometry data with high confidence remains a major bottleneck. We recently introduced a fast and sensitive glycoproteomics search method in our MSFragger search engine, which reports glycopeptides as a combination of a peptide sequence and the mass of the attached glycan. In samples with complex glycosylation patterns, converting this mass to a specific glycan composition is not straightforward, however, as many glycans have similar or identical masses. Here, we have developed a new method for determining the glycan composition of N-linked glycopeptides fragmented by collision or hybrid activation that uses multiple sources of information from the spectrum, including observed glycan B- (oxonium) and Y-type ions and mass and precursor monoisotopic selection errors to discriminate between possible glycan candidates. Combined with false discovery rate estimation for the glycan assignment, we show this method is capable of specifically and sensitively identifying glycans in complex glycopeptide analyses and effectively controls the rate of false glycan assignments. The new method has been incorporated into the PTM-Shepherd modification analysis tool to work directly with the MSFragger glyco search in the FragPipe graphical user interface, providing a complete computational pipeline for annotation of N-glycopeptide spectra with FDR control of both peptide and glycan components that is both sensitive and robust against false identifications.

## Introduction

Glycosylation is one of the most common post-translational modifications of proteins, involved in a vast array of biological processes and implicated in numerous diseases (1–4). Due to the analytical challenges resulting from the heterogeneity of glycosylation, both in sites occupied and glycans present at a given site, analysis of the glycoproteome has generally lagged behind other omics fields (5). Improvements to methods for enriching, separating, and analyzing glycopeptides by mass spectrometry have been accelerating in recent years (5–7), however, resulting in increasingly large and complex glycoproteomics data being generated. Analysis of this data has represented a significant bottleneck in glycoproteomics (8), particularly for proteome-scale analysis of intact glycopeptides. A rapid expansion of software tools is underway in this area, with many new methods capable of this type of analysis being reported recently (9–18).

Statistical control of the results reported by these new tools has fallen behind, however, in large part to the extra challenges of correctly identifying intact glycopeptide spectra (19). As a result, despite the accelerating development of these tools, it remains common practice to manually validate and/or empirically filter search results to remove incorrect glycan composition assignments (20), presenting a major bottleneck for large-scale glycoproteomics studies. Many tools for glycoproteomics data analysis adapt methods from proteomics for glycoproteomics by treating glycans similarly to other post-translational or chemical modifications of peptides. Some search tools provide additional capabilities that can assist in controlling the false discovery rate (FDR) of modified peptides, such as the use of the extended mass model of PeptideProphet (21) to model distinct probabilities for modifications with different masses used with MSFragger (17), or distinguishing between rare and common modifications in Byonic (12). These tools and many others have increasingly been applied to large scale glycoproteomics analyses (14, 18, 22–25) utilizing peptide-focused FDR methods, often in conjunction with a second empirical filtering or manual validation step.

Several studies have pointed out, however, that peptide-focused FDR approaches can fall short for glycoproteomics analyses due to the complexity and heterogeneity of glycans (8, 19, 26, 27). There can be hundreds of different glycans present in an individual glycoproteomics analysis with frequencies that vary over multiple orders of magnitude. Furthermore, N-glycans are comprised of a common core and various extensions, often containing repeating carbohydrate units. There are some residue combinations that are isomeric (e.g., N-glycolyl neuraminic acid (NeuGc) plus fucose has the same atomic composition and exact mass as N-acetyl neuraminic acid (NeuAc) plus hexose), and several more that are very similar in mass to other combinations or to peptide modifications (28–30). Errors in assigning the monoisotopic mass of the precursor, also called “off-by-X” or peakpicking errors, result in a glycan mass that is off by 1 (or several) Da, and there are several additional combinations of common carbohydrate residues or peptide modifications that are very similar in mass to another combination plus such an isotope error (20). Treating glycopeptides as modified peptides and using adapted proteomics methods may thus be sufficient for analyses in which the glycan population is relatively simple and well characterized, and any overlapping glycan masses can be watched for and resolved manually if needed. This manual verification remains a major bottleneck of glycoproteomics analyses, however, and often precludes large-scale analysis of more complex or uncharacterized glycan populations where such overlaps are common.

Several recent methods have been proposed in which a target-decoy analysis is performed specifically on the glycan-matching portion of glycopeptide-spectrum matching to generate a glycan-specific FDR or “glycan FDR” (10, 13, 31). In more recent examples, this has been combined with a “peptide FDR” typical to modern proteomics methods to evaluate the quality of match between the spectrum and both the proposed peptide sequence and glycan composition (9, 11, 32, 33). This approach has the potential to enable automated analysis of glycoproteomics data with complex and uncharacterized glycan populations and remove the manual validation bottleneck. The proposed methods thus far have generally used a “glycan-first” approach, in which possible glycan candidates are first identified by matching the Y-ion series from the spectrum, then the determined glycan mass is subtracted from the observed precursor to determine the peptide mass, and finally the spectrum is searched for matching peptide fragment ions from precursors matching the determined peptide mass. While this method has been shown to be effective at controlling glycan FDR, it is limited to data in which abundant Y-ions are produced, making it challenging to adapt for O-glycoproteomics and potentially its reducing sensitivity when fragmentation conditions are not optimal for producing Y-ions.

Here, we propose a new approach for identifying the glycan component of an N-linked glycopeptide and an associated glycan FDR estimation method with two major differences from the existing methods. We first identify the peptide sequence, using a mass offset-style glyco search from MSFragger (17, 34), then match the mass difference between the peptide sequence mass and the observed precursor mass to candidate glycans to determine the composition. This “peptide-first” approach leverages the well-developed capabilities of modern proteomics methods to solve the peptide portion of glycopeptide identification first, reducing the glycan identification portion to distinguishing between a few glycans that match the mass difference from the determined peptide sequence, rather than distinguishing between the complete search list of up to hundreds of possible glycan compositions. Second, we generate a composite glycan score from a variety of spectral evidence, including Y-ions, oxonium ions, and the observed mass and precursor isotope errors, rather than just Y-ions alone. We demonstrate that by simplifying the problem by matching the peptide first using our MSFragger search tool and then maximizing the glycan-specific information gleaned from spectra, we are able to annotate several-fold more glycopeptide spectra at the same FDR as existing glycan-first methods. Unlike our previous peptide-only FDR approach, we show that this method controls glycan FDR across many search scenarios, including when searching for entrapment glycans known not to be present in the sample, while maintaining the high sensitivity of the MSFragger glyco search. The method has been implemented in the open-source tool PTM-Shepherd (35) v1.2 and has been incorporated into the FragPipe graphical interface and pipeline to provide a complete solution for glycoproteomics analyses.

## Experimental Procedures

### Dataset Details

Raw data were downloaded from Proteome Xchange (36) repositories and converted to mzML with MSConvert (37). The “yeast” dataset from PXD005565 contains glycopeptides enriched by ZIC-HILIC from fission yeast (*Schizosaccharomyces pombe*) analyzed by stepped-energy HCD on an Orbitrap Fusion mass spectrometer (9). The “Riley” dataset from PXD011533 contains glycopeptides enriched by lectin affinity chromatography from mouse brain tissue analyzed by HCD and AI-ETD fragmentation on an Orbitrap Fusion Lumos mass spectrometer (38). The “mouse” 5-tissue dataset contains glycopeptides enriched from mouse brain (PXD005411), kidney (PXD005412), heart (PXD005413), liver (PXD005553), and lung (PXD005555) analyzed by stepped-energy HCD on an Orbitrap Fusion mass spectrometer (9).

### Glycoproteomics Searches and Peptide FDR

Analysis of the data was performed in two parts: first, glycopeptide search using MSFragger’s glyco mode with peptide validation and FDR filtering in Philosopher, and second, glycan assignment and glycan FDR filtering in PTM-Shepherd. MSFragger glyco search, described previously in (17), produces a list of peptide-spectrum matches (PSMs) for both non-glyco and glycopeptides found, in which glycopeptides are matched as a peptide and a delta mass corresponding to the mass of the glycan (Fig. 1A).

**Figure 1.**
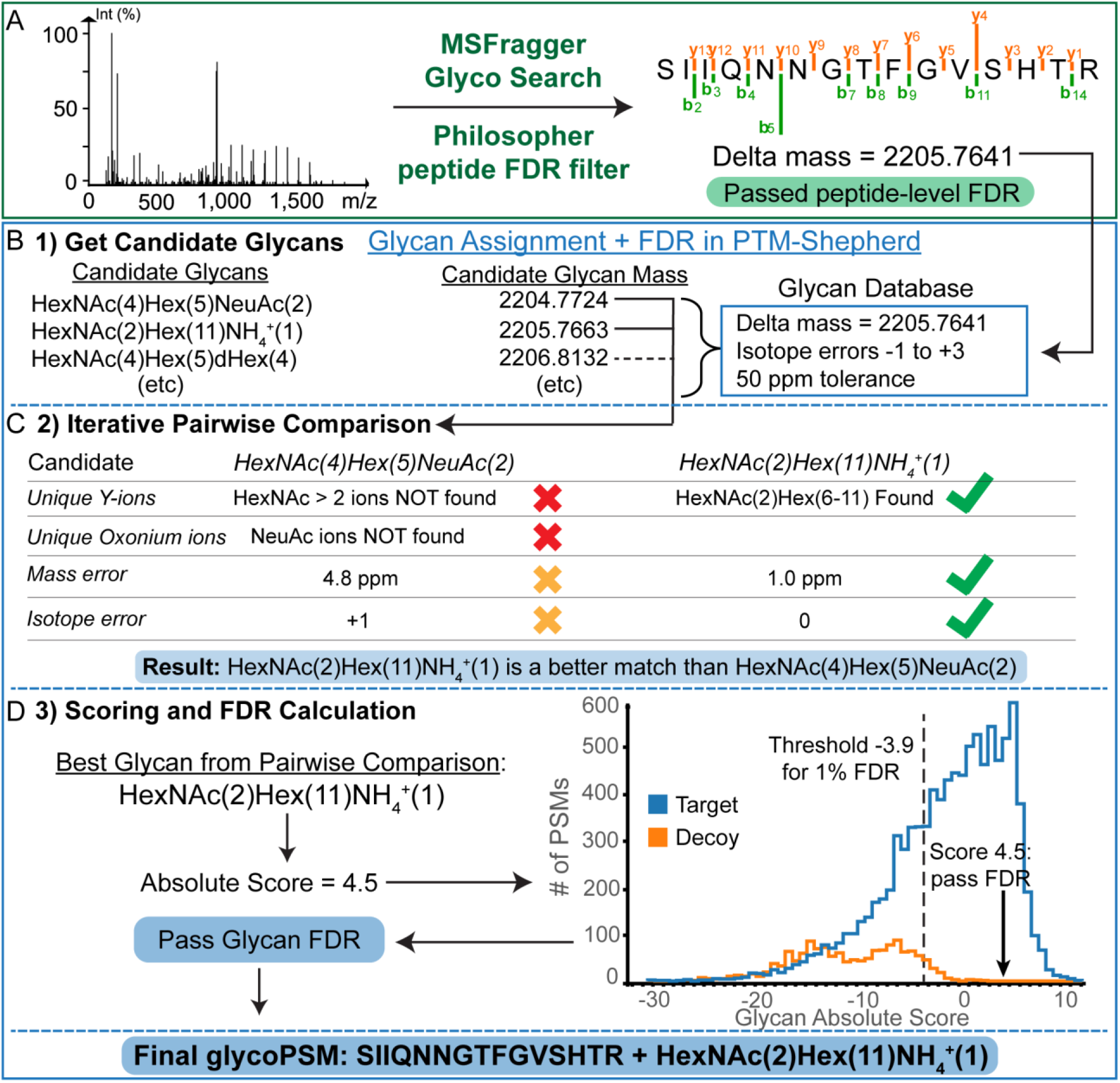
Glycan assignment workflow in PTM-Shepherd. A) Glyco search in MSFragger and peptide FDR filtering in Philosopher annotates a spectrum with a peptide and a delta mass. B) Possible glycan candidates with masses similar to the provided delta mass are gathered from the internal glycan list in PTM-Shepherd. C) Pairwise comparison of all candidates determines the best match to the spectrum based on unique fragment ions for each candidate and mass and isotope errors. Note that mass error is calculated after first correcting any isotope error. Green check marks in a candidate’s column indicate positive evidence for that candidate, red and orange x’s indicate negative or slightly negative evidence, respectively. D) The best candidate is re-scored to generate the absolute score and FDR is computed using the score distribution of target and decoy glycans.

“Yeast” data from PXD005565 was converted to mzML format using MSConvert (37) with vendor peakpicking and searched against a combined yeast and mouse proteome database with common contaminant proteins and decoys appended in Philosopher (downloaded 02/08/2021, 22,307 non-decoy entries). Searches were performed against several glycan lists: a “yeast” list containing only glycans with HexNAc and Hex residues and up to 1 fucose or ammonium adduct (186 unique masses), a “mouse” list that included glycans with HexNAc, Hex, dHex (e.g., fucose), and NeuAc-containing compositions (182 unique masses), and large mammalian N-glycan list (“full list”), equivalent to the “pGlyco-N-mouse-large” list (containing 1670 unique compositions). One ammonium adduct was allowed, increasing the total number of unique mass offsets searched in MSFragger to 2325 (redundant masses (rounded to 2 decimal places) were removed). This full list was the same as used in the analysis of the yeast dataset in the manuscripts describing the pGlyco2 (9) and pGlyco3 (11) software packages. The full glycan lists can be found in Supplementary Data 1. MSFragger (v3.2) searches were performed against the combined yeast/mouse database, allowing 2 missed cleavages by Trypsin, fixed carbamidomethylation of Cys, and variable Met oxidation and protein N-terminal acetylation, in n-glycan mode with precursor and fragment mass tolerances of 20 and 10 ppm, respectively, precursor isotope error correction enabled using the built-in correction algorithm, *b*, *y*, *b* + HexNAc, *y* + HexNAc, and Y-ions considered, and oxonium ion filtering enabled (minimum 10% summed oxonium ion intensity relative to the base peak of the spectrum to search for glycopeptides) with default oxonium ion masses. Following MSFragger search, Philosopher (v4.0.0) (39) was used to perform peptide FDR filtering. PeptideProphet (21) with extended mass model (mass width 4000) was used to model PSM probabilities in semi-parametric mode with n-glycan motif modeling enabled and cLevel set to 0. ProteinProphet (40) protein inference was performed with default parameters except maxppmdiff set to 20000000 to prevent exclusion of glycopeptides with large delta masses. The final PSMs, peptides, and proteins were filtered to 1% PSM and protein FDR, using a sequential filtering step to remove PSMs and peptides from proteins that did not pass FDR.

Results from a pGlyco3 primary search (without ammonium adducts) of the yeast dataset shown in Figure 2 performed as part of (11) were downloaded from the Massive repository MSV000086771. pGlyco3 (build 20210615) was used to perform an equivalent search with ammonium adducts. The same combined yeast/mouse proteome database as the MSFragger/PTM-Shepherd search (above) was used, and the “p-Glyco-N-mouse-Large” glycan database. Raw files were parsed with pParse using default parameters and peakpicking. Parameters for protein digestion, peptide modifications, and mass tolerances were set to the same values as in the MSFragger search. One “aH” variable modification was allowed on glycans (ammonium adduct). Default FDR options were employed (1% peptide and glycan) using the built-in peptide FDR method.

**Figure 2.**
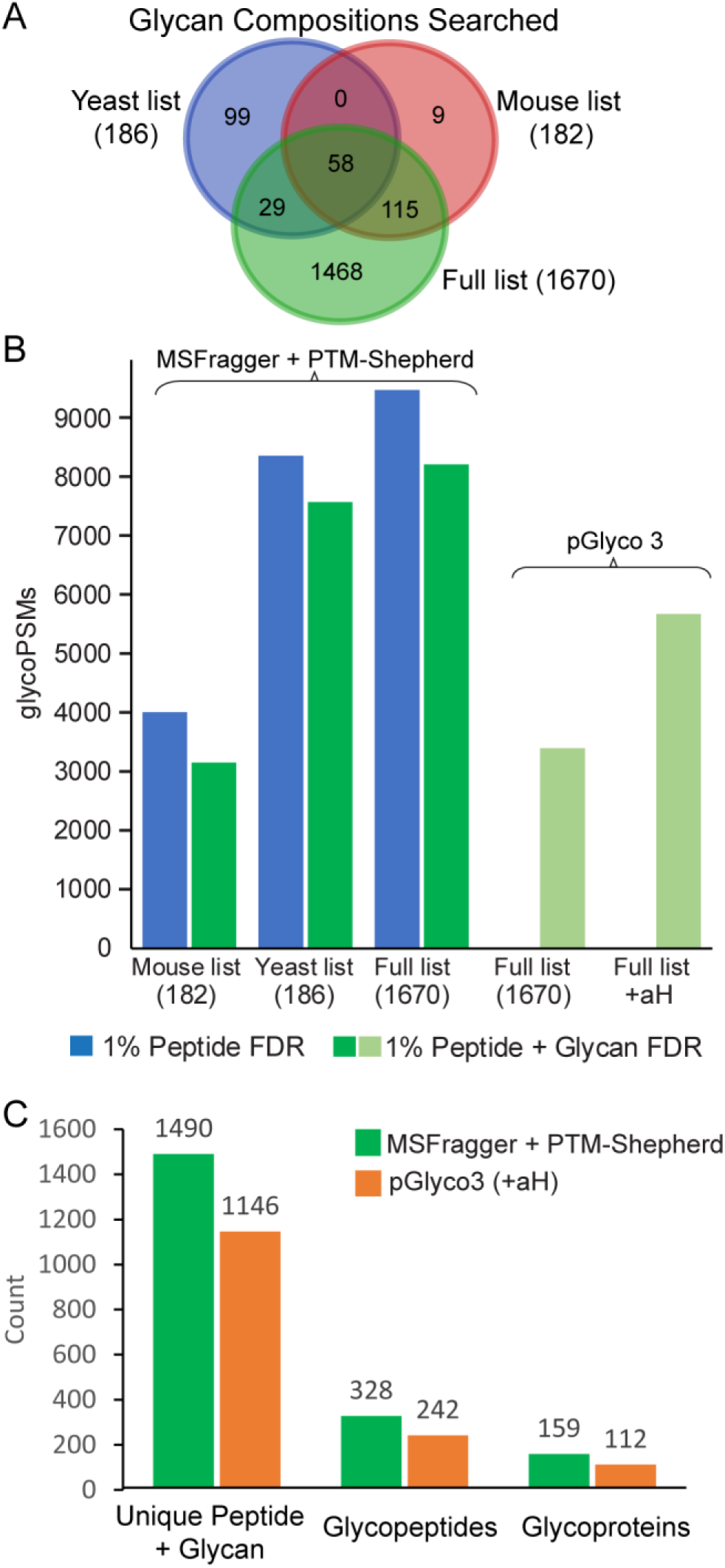
Results of entrapment searches of yeast dataset. A) Venn diagram of the three glycan lists used in searches showing the overlap in compositions between searches. B) GlycoPSMs annotated at 1% peptide FDR (blue) and combined 1% peptide and glycan FDR (green) for MSFragger searches of three different glycan lists with PTM-Shepherd glycan FDR filtering. Comparison with pGlyco 3 reported results for the same peptide database, glycan list, and FDR levels is at right either with or without allowing 1 ammonium adduct per glycan (encoded by pGlyco3 as “aH” residues). C) Unique glycoproteins, glycopeptide sequences, and glycan/peptide combinations (each glycan composition on each unique peptide sequence counts as a separate entry) detected in MSFragger + PTM-Shepherd (green) and pGlyco3 (orange) results at 1% peptide and glycan FDR.

The “Riley” dataset from PXD011533 was searched and FDR filtered with the same method and parameters except for the following differences. HCD and AI-ETD scans were extracted to separate mzML files with vendor peakpicking in MSConvert and searched separately before results were combined for validation and FDR filtering. Spectra were searched against mouse glycoprotein-focused database from Riley et. al. (38) with decoys appended in Philosopher (3,574 entries non-decoy entries), allowing up to 3 missed cleavages, and the mouse glycan list (182 unique masses). AI-ETD search considered *b*, *y*, *c*, *z*, and Y-ions and reduced required oxonium ion intensity to 2.5%.

“Mouse” data from PXD005411, PXD005412, PXD005413, PXD005553, PXD005555 were converted to mzML with MSConvert using vendor peakpicking and searched against a full reviewed mouse proteome with common contaminant proteins and decoys appended in Philosopher (downloaded 09/24/2019, 17,019 non-decoy entries). The same “full list” of glycans was used as in the yeast analysis, equivalent to the “pGlyco-N-Mouse-Large” glycan list with one ammonium adduct allowed. All other MSFragger parameters were the same as in the yeast searches. An entrapment search of the mouse data was also performed, with 248 entrapment glycan compositions added to the MSFragger search and PTM-Shepherd glycan list (see Supplementary Data 1 for the list of entrapment glycans).

Comparative searches of the mouse data were performed with pGlyco3 (build 20210615) using the same mouse protein database and glycan list with 1 “aH” variable glycan modification (ammonium adduct) allowed. Raw data was read with pParse using default parameters and peakpicking. Variable peptide modifications were set to the same as MSFragger search (protein N-terminal acetylation and Met oxidation) and precursor and fragment mass tolerances were set to 20 and 10 ppm, respectively, as in the MSFragger search. Default FDR options were used (1% peptide and glycan FDR) using the built-in peptide FDR method.

### Converting Delta Mass to Glycan Composition in PTM-Shepherd

PTM-Shepherd (35) reads PSM results, including the identified peptide and delta mass, from the output of MSFragger and Philosopher. Determining the identity of the glycan from the delta mass begins by determining possible glycans with intact masses near the observed delta mass from a provided list of glycan compositions or PTM-Shepherd’s internal glycan list (Fig. 1B). This internal list is constructed from reported N- and O- glycans from several glycoproteomics analyses and databases with the intent of including most known mammalian glycan compositions so that it can be used without modification for a range of glycoproteomics analyses(the list can be found in Supplementary Data 2). Custom lists of glycan compositions can also be supplied by the user and should be used when analyzing samples with glycosylation that differs markedly from mammalian N-glycosylation. If specified by the user, adducted forms of glycans can be considered, e.g., the replacement of a proton by an ammonium adduct, which does not change the expected fragment ions, as the noncovalent adduct is not expected to be retained. The current version of the method supports only peptides with a single glycosylation event, though glycopeptides with multiple potential glycosylation sites are not explicitly excluded from analysis and can thus be matched if only one site is in fact glycosylated.

To allow for FDR control, a decoy glycan is appended to the list for each target (and each target-adduct, if specified). A decoy glycan candidate is generated by shifting the intact mass of a target glycan by a random value within the provided glycan mass error tolerance and assigning a randomly chosen isotope error from the set of such errors being considered in the analysis. This allows us to distinguish decoy glycans from targets on the basis of their different mass and isotope error distributions in addition to Y and oxonium ions, while ensuring that decoys are not being shifted to masses that would exclude them from consideration alongside their target glycans. A decoy glycan candidate has the same nominal composition as the target glycan from which it was generated, but its fragment (Y and oxonium) ions are also each randomly shifted by a unique value between 1 – 20 Da. As a result, the decoy has the same number of fragment ions of each type as its corresponding target, but with randomly shifted masses.

Target and decoy glycans within a user-specified tolerance of the observed delta mass (50 ppm used for all analyses here), and with allowed precursor isotope errors (−1, 0, +1, +2, or +3 for all analyses here) are considered as possible candidates (Fig. 1B). The observed isotope error for a candidate is determined by subtracting the candidate mass from the observed mass and rounding to the nearest integer. The resulting isotope error mass, which is the determined isotope integer error times an average peptide isotope spacing of 1.00235 Da, is removed from the candidate mass prior to computing the mass error score.

For each spectrum with a mass shift potentially corresponding to a glycan, pairwise comparisons are then performed to determine the best candidate glycan from the list of possible candidates within tolerance of the observed delta mass from the available evidence (Fig. 1C). Starting from two arbitrary candidates, the current best candidate is compared to other candidates until all candidates have been considered, with current best candidate being updated any time a compared candidate generates a higher score. Candidates are compared using four components that are ultimately combined into a single score: Y-ions, oxonium ions, mass error, and isotope error. For Y- and oxonium ions, scoring is based on ions that are unique to one candidate or the other; ions that can be generated by both candidates or neither candidate have no impact on the score. The pairwise scoring function is based on summed log likelihood estimation of the impact of each piece of evidence (each Y- or oxonium ion and the observed mass and isotope errors) on the likelihood of the spectrum representing one of the candidates rather than the other (Eqn. 1–3). For Y- and oxonium ions unique to one of the candidates, an empirically determined probability ratio is used to express the effect of observing the ion on the likelihood of the spectrum representing that candidate rather than the other. Currently, Y-ions are divided into 2 categories based on whether or not they contain Fucose, and all ions within each category are given the same score. The Y-ion score can thus be expressed as a constant α, representing the log of the hit or miss probability ratio, times the number of unique Y-ions found or not found for each candidate for each of non-Fucose and Fucose containing Y-ion sets (Eqn. 2). Because the number of Y-ions observed tends to be less than the number of possible Y-ions, particularly for larger glycans, Y-ion counts are square root normalized to avoid over-penalizing large glycans with more possible Y-ions than are typically observed. Probability ratios for oxonium ions are encoded separately for several composition categories (currently NeuAc/NeuGc (41–43), Fucose (44, 45), phosphate (46), and sulfate (47, 48) are supported based on our empirical observations and oxonium ions reported elsewhere (49, 50), resulting in the ratios shown in Supplementary Table S1. The oxonium ion score is thus computed similarly to the Y-ion score, except with potentially different probability ratios for each ion type and without square-root normalization of ion counts (Eqn. 3). Following the observation that oxonium ions resulting from co-fragmentation of glycopeptides had a negative impact on assignment quality, an intensity weighting factor was added to the oxonium fragment score. The hit probability ratio is multiplied by the ratio of observed divided by expected intensity, such that low intensity oxonium ions will result in a smaller increase in score than more intense ones. The adjustment is capped so that finding a hit with extremely low intensity cannot negatively impact the score: in these cases, the low intensity hit results in no change to the score. The mass error score is the log of the ratio of observed mass errors for candidates 1 and 2, such that a lower mass error for candidate 1 than candidate 2 results in a positive score, with a weight factor β (set to 1 by default). The isotope error score is the log of the probability ratios (α) for the observed isotope errors of each candidate (see Supplementary Table S2 for the values used). The equation for the pairwise score of glycan candidate 1 vs candidate 2 is thus:

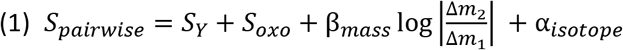

where scores for Y- and oxonium ions (*S*_*Y*_ and *S*_*oxo*_) are defined below.

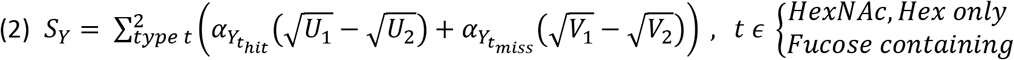

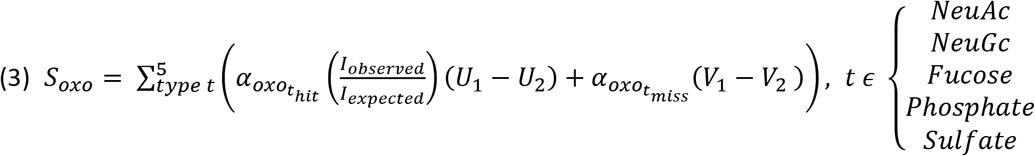

U_1_ is the number of unique fragment ions (Y-ions in equation 2 or oxonium ions in equation 3) to glycan candidate 1 found in the spectrum (unique hits) and V_1_ is the number of unique fragment ions from glycan 1 not found in the spectrum (unique misses), and U_2_ and V_2_ are analogous for glycan candidate 2. I is the intensity of a fragment ion observed in the spectrum or “expected” (provided as a parameter). The ratio of observed to expected intensity has a minimum set such that α cannot be negative for very low observed intensities; in these cases, α is set to 0. For Y- and oxonium ions, probability ratios for hits are always greater than 1 (leading to α > 0) and always less than 1 for misses (α < 0), so that hits for candidate 1 increase the score and misses decrease it, while hits for candidate 2 decrease the score and misses increase it. Probability ratios and expected intensities used for each ion type and isotope error can be found in Supplementary Tables S1-2.

After the best candidate glycan is determined by pairwise comparison for each PSM, FDR estimation is performed. Because the candidate’s score in pairwise comparison depends both on the candidate and the identity of the next-best candidate, it is not optimal for distinguishing between targets and decoys. Instead, an “absolute” score is computed for the top-ranked glycan from pairwise comparison for each PSM (Eqn. 4). This absolute score is similar to the pairwise comparison score but treats all fragment ions from the best candidate glycan as unique and compares against typical mass and isotope errors rather than those of another candidate. Intuitively, the absolute score can be viewed as the total weight of evidence for and against the best candidate.

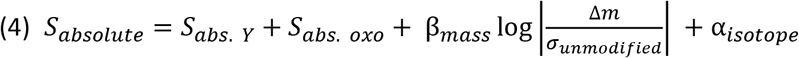

Because there is no second candidate to compare against, probability ratios for the mass and isotope errors of the candidate are compared to typical values. The average mass error of all unmodified peptides in the analysis (*σ*_*unmodified*_) is used as a typical value for mass error and no isotope error is used as the typical value for isotope error. Y- and oxonium ion scores are computed in the same fashion as in the pairwise score but without a second candidate, treating all possible fragment ions from the chosen candidate as unique (and thus contributing to the score):

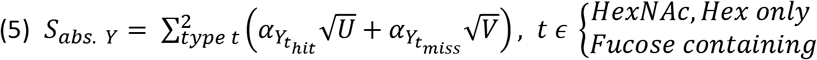

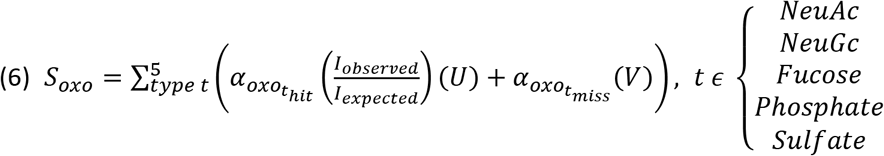

Where U is the number of theoretical fragment ions from the candidate found in the spectrum, analogous to the unique hits from the pairwise score, and V is the number of theoretical fragment ions not found, analogous to the unique misses from the pairwise score. I is the intensity of the fragment (expected and observed) as in the pairwise scoring, and has the same minimum value to prevent α from turning negative for very low observed intensities.

FDR is computed by collecting absolute scores of all target and decoy best candidates and determining the score threshold necessary to achieve the desired decoy-target ratio (Fig. 1D). FDR is computed as the ratio of decoys to targets at a given score value. Results are reported in the PSM results table with the q-value and identity of the matched candidate. By default, PSMs that do not pass glycan FDR are still reported but can be removed by filtering by the appropriate q-value cutoff (e.g., less than 0.01 for 1% glycan FDR). PSMs in which a decoy glycan was assigned instead have the best target glycan reported but with a q-value of 1. An option is available to instead print decoy glycan assignments directly for diagnostics.

All datasets tested used the default probability ratios for fragment ions and mass and isotope errors displayed in Supplementary Tables S1-2. Glycans with noncovalent ammonium (NH_4_^+^) adduct(s) were appended to the internal PTM-Shepherd database in the “yeast” and “mouse” analyses as well after testing revealed high prevalence of these adducts. The “Riley” dataset was searched with no adducts.

### Experimental Design and Statistical Rationale

The yeast dataset from PXD005565 contains a single sample analyzed in technical triplicate. The Riley dataset from PXD011533 also contains a single sample analyzed in technical triplicate (and fractionated into 12 fractions per replicate). The mouse dataset contains a single sample from each tissue type analyzed in 5 technical replicates. All data analyzed is being used for method development and no biological conclusions are drawn. The output of MSFragger search is deterministic, as such, each MSFragger search was performed only once. PTM-Shepherd glycan assignment scoring is deterministic, but the method of generating decoys by randomly shifting the masses of decoy glycans and their fragments would result in different decoy scoring, and thus different FDR thresholds, in replicate runs. As generating different results for repeated analyses of the same input data and parameters is not desirable, we have opted to fix the random seed used to generate decoys, as is done in other software tools employing this decoy generation strategy. Thus, the same decoys will always be generated for the same input raw data and parameters but will be different (randomly) if the input raw data, glycan database, or search parameters are changed.

## Results and Discussion

The method presented here identifies the glycan composition represented by the mass shift reported by MSFragger glyco search and uses the target-decoy approach to enable FDR filtering of the identified glycans to a defined confidence level. Unlike many PTMs, the heterogeneity of glycosylation and abundance of monosaccharide combinations with very similar or identical masses can make it challenging to determine the glycan composition represented by a given mass shift, particularly in the analysis of complex glycosylation profiles. Glycan compositions can also correspond to multiple structures resulting from various connection points between monosaccharides and different branching, however, directly determining glycan structure from glycopeptide fragmentation data is extremely challenging and typically requires specialized MS methods. All results reported here aim to confidently identify a glycan composition, i.e., the identity and count of all monosaccharide classes (e.g., hexose rather than specific monosaccharides like glucose) present, from typical glycoproteomics MS data without implying specific connectivity or branching information. To assess the performance of the glycan assignment method in PTM-Shepherd, we first turned to a well-characterized fission yeast dataset (PXD005565) (9). The fission yeast analyzed therein have a relatively simple glycosylation profile, with the vast majority of glycans consisting of HexNAc(2)Hex(n) structures, with n ranging from 4 to approximately 20. Following the example set by several other studies that use this data for benchmarking (9, 11, 32, 51), we performed an entrapment analysis by searching the yeast data against both yeast and mouse proteomes and glycomes to evaluate the accuracy of both peptide and glycan assignment in the presence of peptides and glycans not expected to be present in the sample. MSFragger searches were performed with three different glycan mass lists: a “yeast-only” list, a small “mouse” list, and a “full” list equivalent to the large mouse glycan list used to perform an entrapment search in this data by pGlyco3 (11) (Fig. 2A, Supplementary Data 1). In all cases, the number of mouse peptides matched was well controlled by the MSFragger pipeline (Supplementary Table S5). The output of the MSFragger search, a peptide and mass shift for each spectrum, was then analyzed using PTM-Shepherd to assign the glycan corresponding to the mass shift for each PSM using the “full” glycan list containing both yeast and entrapment glycans in all cases.

The search results from the three different glycan lists illustrate several important factors in glycan assignment and the performance of our method in a variety of situations. Not all glycans in the “mouse” list are entrapment glycans, i.e., there is some overlap with the yeast list (Fig. 2A), and the search results from MSFragger reflect this, matching roughly half the number of glycoPSMs as when searching with the yeast glycan list. If glycan identities are assumed by naively matching the delta masses observed to the closest mass in the mouse glycan list, a large number of erroneous identifications result (i.e., glycans containing NeuAc, which is not present in fission yeast) (Supplementary Table S4). However, when assigning glycans in PTM-Shepherd (using the full internal N-glycan list containing additional entrapment glycans, not just the 182 mouse glycans searched in MSFragger) at 1% glycan FDR, the rate of erroneous assignment is well controlled, with 0.5% of glycoPSMs matching entrapment glycans out of 3,170 glycoPSMs reported after glycan FDR filtering (Table 1, Fig. 2B). When searching with yeast-specific glycans, MSFragger matches more than twice as many potential glycoPSMs, and because the list of glycans searched closely matches the actual glycan population, glycan FDR filtering in PTM-Shepherd removes a smaller fraction of the matched spectra, yielding 7,598 total glycoPSMs at 1% peptide and 1% glycan FDR (Table 1). Expanding the search space to the general glycan list, with nearly 10 times as many glycan masses considered, results in a moderate further increase in potential glycoPSMs from MSFragger, resulting in 8,234 glycoPSMs at 1% peptide and glycan FDR (Table 1). Crucially, despite the nearly 10-fold increase in glycan search space, the number of PSMs matched to entrapment glycans not present in fission yeast (i.e., glycans containing NeuAc, NeuGc, Fucose) remains extremely well controlled, with only 2 such entrapment PSMs reported in any category after FDR filtering (Table 1). The mouse glycan list, which contains many glycans not present in the data and is also missing many glycans that are actually present has a large number of incorrectly matched spectra that are removed by filtering. The searches with yeast-specific and full lists, which contain nearly all glycans that actually present, have a moderate number of spectra removed by filtering, lower in the yeast-specific list search. The majority of these removals are due to low quality fragmentation failing to provide sufficient evidence for the correct composition. There were a few cases where an incorrect peptide assignment or missing peptide modification resulted in a low-quality glycan match that was removed by filtering. Some peptides with two potential N-glycosylation sites were assigned glycan compositions that appeared to be the combination of two glycans, which generated very low glycan assignment scores and were also removed. Unlike the peptide-only FDR control of the original MSFragger glyco search, which provided reasonable assignments only when the searched glycans are well matched to the actual glycans present in the data, the method presented here was able to control the rate of entrapment matches regardless of the glycan list being searched.

**Table 1.**
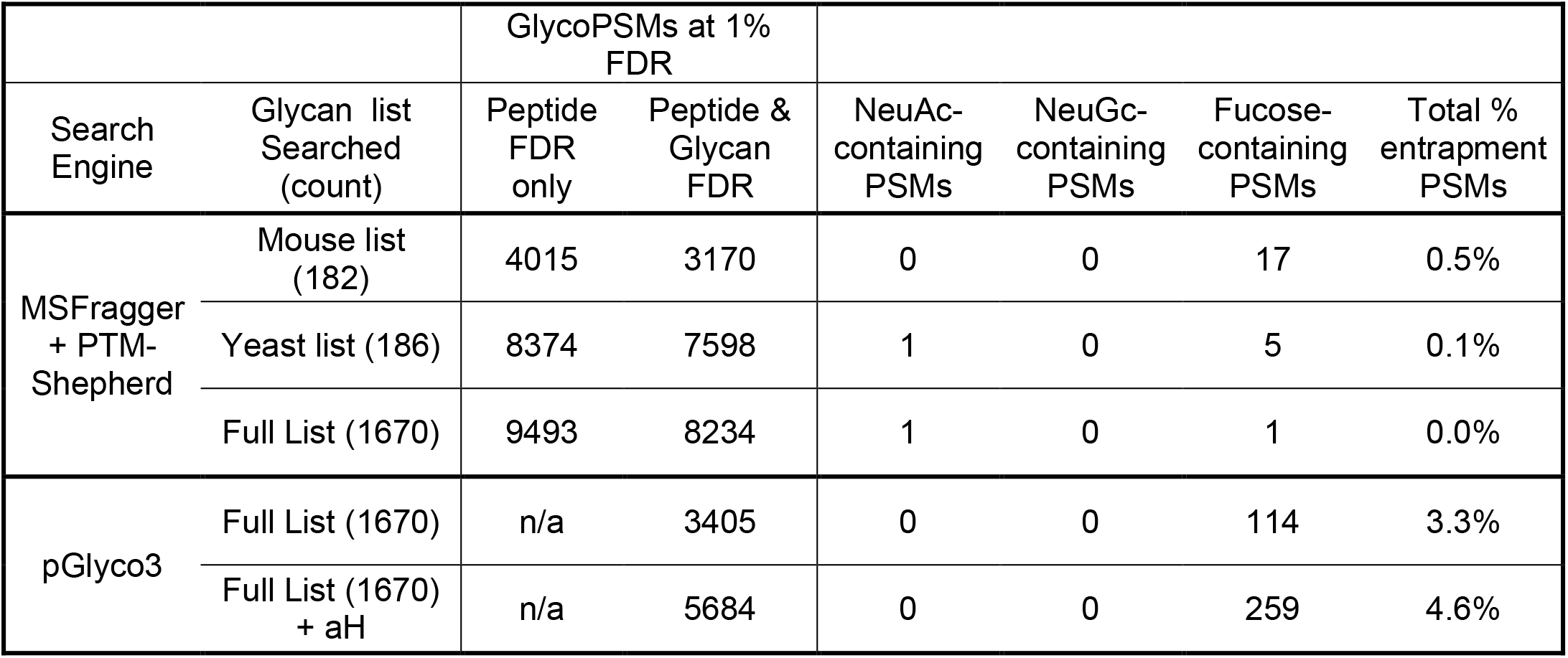
Results from entrapment searches of yeast data at 1% FDR. GlycoPSMs annotated at 1% peptide and combined 1% peptide and glycan FDR are shown for searches of three different glycan lists, along with the number of PSMs annotated with various entrapment glycans for each search. Total percentage of glycoPSMs matched to entrapment glycans is shown at right. Top three rows of results are from MSFragger and PTM-Shepherd (“PTM-S”); the bottom two rows contain results from pGlyco3 (11) using same full list of glycans. The bottommost row allows ammonium adduction of glycans (what pGlyco3 calls an “aH” glycan modification).

Another state-of-the-art software package, pGlyco3, recently analyzed this same data (11), allowing for a direct comparison. At the same nominal FDR (1% peptide and 1% glycan), pGlyco3 reported 3,405 glycoPSMs when not considering ammonium adducts or 5,684 glycoPSMs when allowing an ammonium adduct, compared to the 8,234 from MSFragger/PTM-Shepherd when searching the same peptide database and glycan list and allowing an ammonium adduct. pGlyco3 reported no PSMs containing NeuAc or NeuGc, compared to 1 PSM with NeuAc or NeuGc reported by MSFragger/PTM-Shepherd, indicating good FDR control for sialylated glycans by both tools. However, pGlyco3 also reported 114 PSMs (3.3% of all glycoPSMs) containing Fucose, or, when allowing ammonium adduction, 259 PSMs with Fucose (4.6%), which are unlikely to be present in the yeast dataset. MSFragger/PTM-Shepherd reported only 1 such PSM, indicating much improved FDR control for these glycans (Table 1). Our method is thus able to annotate nearly 50% more glycoPSMs while also providing robust FDR control in more types of glycans than pGlyco3.. We attribute the dramatic difference in performance between our method and that of pGlyco3 primarily to two factors, the peptide-first search approach resulting in many more possible glycoPSMs being considered in glycan assignment and incorporation of multiple types of information to glycan scoring in PTM-Shepherd. In particular we found that inclusion of a weak oxonium ion filter for Fucosylated glycans (in addition to the Y-ions considered by both our method and pGlyco3) provided greatly improved control of erroneous matches.

Additional tests confirm the robustness and sensitivity of the glycan FDR estimation in PTM-Shepherd. Results from searches with various glycan FDRs (1%, 0.5%, and 0.1%) are shown in Table 2. At 0.1% glycan FDR, no entrapment glycoPSMs are reported at all, but over 6,700 glycoPSMs are still obtained, more than pGlyco3 at 1% glycan FDR. At each FDR tested, the proportion of entrapment glycans assigned remains far below the assigned FDR. While proportion of entrapment glycans matched is necessarily an underestimation of the true FDR (i.e., just because a yeast glycan is matched does not necessarily mean that it was correctly assigned to a given spectrum), consistently observing almost no entrapment glycans implies robust FDR control. Finally, we tested our method without prior peptide FDR filtering, providing over 33,000 potential glycopeptide spectra from the MSFragger search to PTM-Shepherd instead of only the 9,493 glycopeptide spectra that passed peptide FDR. These additional ∼23,000 spectra with low-confidence peptide assignments represent an increased challenge for glycan FDR filtering, as any incorrectly assigned peptide sequences may have an incorrect delta mass used to determine possible glycan candidates. At each glycan FDR level tested, however, the proportion of non-yeast glycans remained well below the given FDR level in all cases (Table 3), demonstrating the robustness of our glycan FDR approach even when analyzing spectra without controlling the rate of false peptide matches.

**Table 2.** Comparison of search results of yeast data with various glycan FDR levels. Full (1670) glycan list was used for the MSFragger search, resulting in 9,493 potential glycoPSMs passing 1% peptide FDR. Entrapment glycans matched are shown at right for each search, with the total percentage of PSMs matched to entrapment glycans at far right.

| Glycan FDR | 1% Peptide + Glycan FDR glycoPSMs | NeuAc-containing PSMs | NeuGc-containing PSMs | Fucose-containing PSMs | % Entrapment glycans |
| --- | --- | --- | --- | --- | --- |
| 1% | 8234 | 1 | 0 | 1 | 0.0% |
| 0.5% | 8022 | 1 | 0 | 1 | 0.0% |
| 0.1% | 6741 | 0 | 0 | 0 | 0.0% |

**Table 3.** Summary of yeast search results without prior peptide FDR filtering at various glycan FDR levels. Full (1670) glycan list was used for the MSFragger search. In total, 33,450 potential glycoPSMs were supplied to PTM-Shepherd with no peptide FDR filter. PSMs matched to entrapment glycans are shown for each FDR level along with the total percentage of PSMs matched to entrapment glycans.

| Glycan FDR | Glycan FDR only glycoPSMs | NeuAc-containing PSMs | NeuGc-containing PSMs | Fucose-containing PSMs | % Entrapment Glycans |
| --- | --- | --- | --- | --- | --- |
| 1% | 14052 | 1 | 0 | 50 | 0.4% |
| 0.5% | 11291 | 1 | 0 | 13 | 0.1% |
| 0.1% | 5271 | 0 | 0 | 0 | 0.0% |

Compared to existing glycan FDR estimation approaches that rely only or primarily on the Y-ion series, our method uses several additional components to evaluate potential glycan candidates. To evaluate the individual contribution of each of these components, we performed the assignment and FDR procedure sequentially with a single score component removed and assessed the number of PSMs passing glycan FDR and any changes in the entrapment rates of different glycan types (Fig. 3). Of the 4 components, isotope error provided the least valuable contribution, indicated by its removal not substantially changing the number of glycoPSMs passing FDR and only slightly increasing the number entrapment glycans matched (Fig. 3). This is perhaps unsurprising given that MSFragger was set to attempt to correct any errors in monoisotopic peak selection prior to PTM-Shepherd analysis. In cases where such correction is not performed, the isotope error component may provide greater benefit. Removal of the oxonium ion score also resulted in only a small change in the number of PSMs reported, but caused uncontrolled matching to compositions containing sialic acids and, to a lesser extent, fucose . Since the sialic acids are typically dissociated from the glycan in the HCD fragmentation employed in this dataset, they generally do not affect the Y-ions produced and are thus reliant on oxonium ion scores for appropriate scoring and filtering. Removal of the mass error score resulted in a moderate decrease in the number of glycoPSMs matched, indicating that it provides a valuable contribution in distinguishing true matches, but did not cause an increase in entrapment glycans matched. Finally, removal of Y-ions resulted in the largest decrease in glycoPSMs, indicating the central role they play in obtaining confident matches. These results illustrate the importance of including multiple sources of information when evaluating glycans, as Y-ions, oxonium ions, and mass error all substantially improved the ability to distinguish between glycan compositions. The relative contributions of the categories may vary in different analyses, however, as a result of different fragmentation methods or settings changing the likelihood of observing different types of fragment ions, or different instruments or instrument settings changing the distribution of mass and isotope errors compared to the data analyzed here, for example.

**Figure 3.**
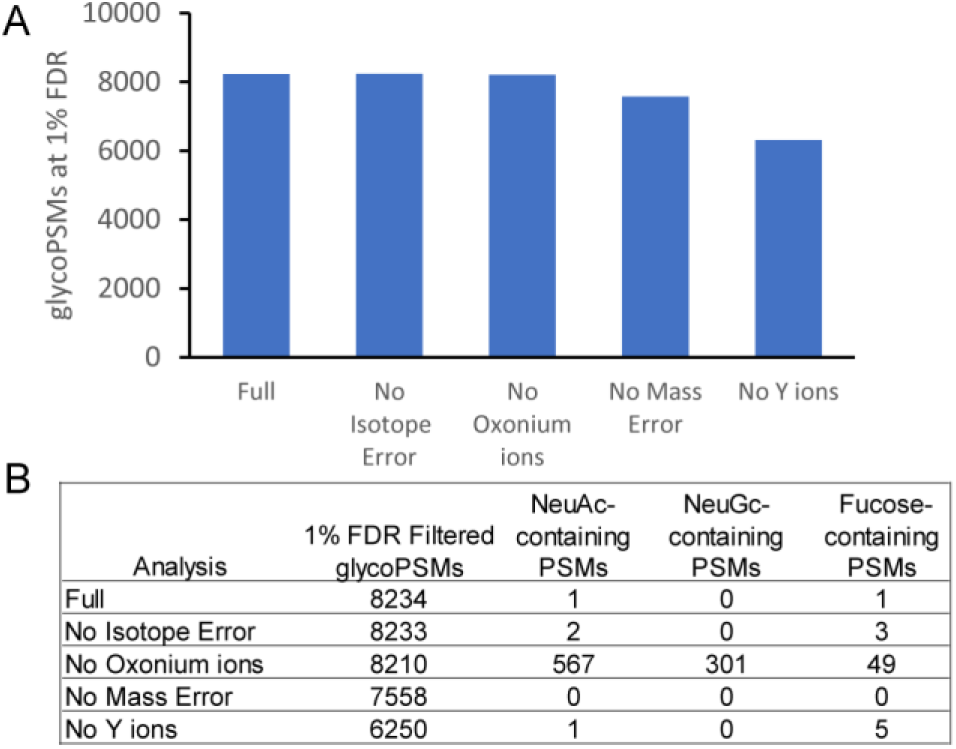
Impact of individual score components on glycan assignment performance. A) glycoPSMs passing 1% glycan (and peptide) FDR with full score (all components) or one component removed. B) Table of total glycoPSMs and entrapment glycoPSMs of various types for each analysis presented in A.

Having validated our glycan assignment and FDR methods with entrapment searches, we set out to characterize the performance of our method in more typical, non-entrapment, glycoproteomics data. We re-analyzed data from Riley *et al*. (38) to compare with our previous analysis by MSFragger-Glyco (17). The original analysis, performed with Byonic (12), identified 24,099 glycoPSMs after applying Byonic’s peptide FDR filtering and additional empirically-determined score filters. Applying the same mouse glycoprotein database and glycan list, MSFragger identified 45,318 glycoPSMs at 1% peptide FDR, of which 44,781 glycoPSMs passed 1% glycan FDR in PTM-Shepherd (Fig. 4A). Unlike the entrapment searches of fission yeast, the glycan FDR filtering removed relatively few PSMs, indicating the high quality of the initial matches from MSFragger when searching only for glycopeptides that are likely to be present in the data. However, the glycan assignment method is still critical to distinguish the correct glycan out of multiple possibilities, even when only expected glycans are included in the search. The additional glycoPSMs annotated result in identification of additional glycopeptides and glycoproteins, as we have previously noted, and additional high confidence pairings of peptide sequences with specific glycan compositions (Fig. 4B). We find similarly small proportions of glycan compositions containing fucose and sialic acids as in the original analysis of Riley *et al.* Mouse brain tissue has been observed to have a high proportion of oligomannose-type glycans (9, 52) (which do not contain fucose or sialic acids), and the Concanavalin A lectin enrichment performed specifically enriches for oligomannose-type glycans over others, both of which lend support to our assignment of the majority of glycoPSMs to compositions containing only HexNAc and Hex-type residues. While our method assigned different glycans than Byonic to several hundred glycoPSMs, the differences were largely between similar compositions that differ by isotope errors. For example, our method frequently assigned the composition HexNAc(2)Hex(8) to spectra that Byonic assigned as HexNAc(6)Hex(3), often when the precursor was misassigned as the +2 isotope peak, as the two compositions differ in mass by 2.05 Da (Supplementary Table S6). Comparisons of assigned glycans are necessarily limited, however, as Byonic does not perform a glycan-specific FDR calculation, controlling peptide FDR only. While we did not perform a deliberate entrapment search on this data, two alternative methods for confirming our glycan assignments are available since mouse brain tissue is not expected to contain NeuGc or sulfated glycans, and the dataset contains paired HCD and AI-ETD scans of the same precursor. The MSFragger search performed included only 182 glycan compositions, but for the PTM-Shepherd analysis we used a large mammalian N- and O-glycan list that included NeuGc and sulfate-containing glycans (Supplementary Data 1), many of which have similar or identical masses to the glycans included in the MSFragger search. Less than 0.1% of all glycoPSMs were matched to either of these unexpected residue types (Fig. 4C), indicating good FDR control for this data. In addition, the paired HCD and AI-ETD scans should have same peptide and glycan assigned, since they are of the same glycopeptide precursor fragmented using different activation methods. Of the 28,970 glycoPSMs that had a paired scan that passed both peptide and glycan FDR, 99.4% were independently assigned to the same peptide and glycan by our method (Fig. 4D), as would be expected given a 1% peptide and glycan FDR. Notably, while our empirical fragment ion probability ratios were developed using HCD data, performance in glycan assignment was comparable in the hybrid AI-ETD data included here.

**Figure 4.**
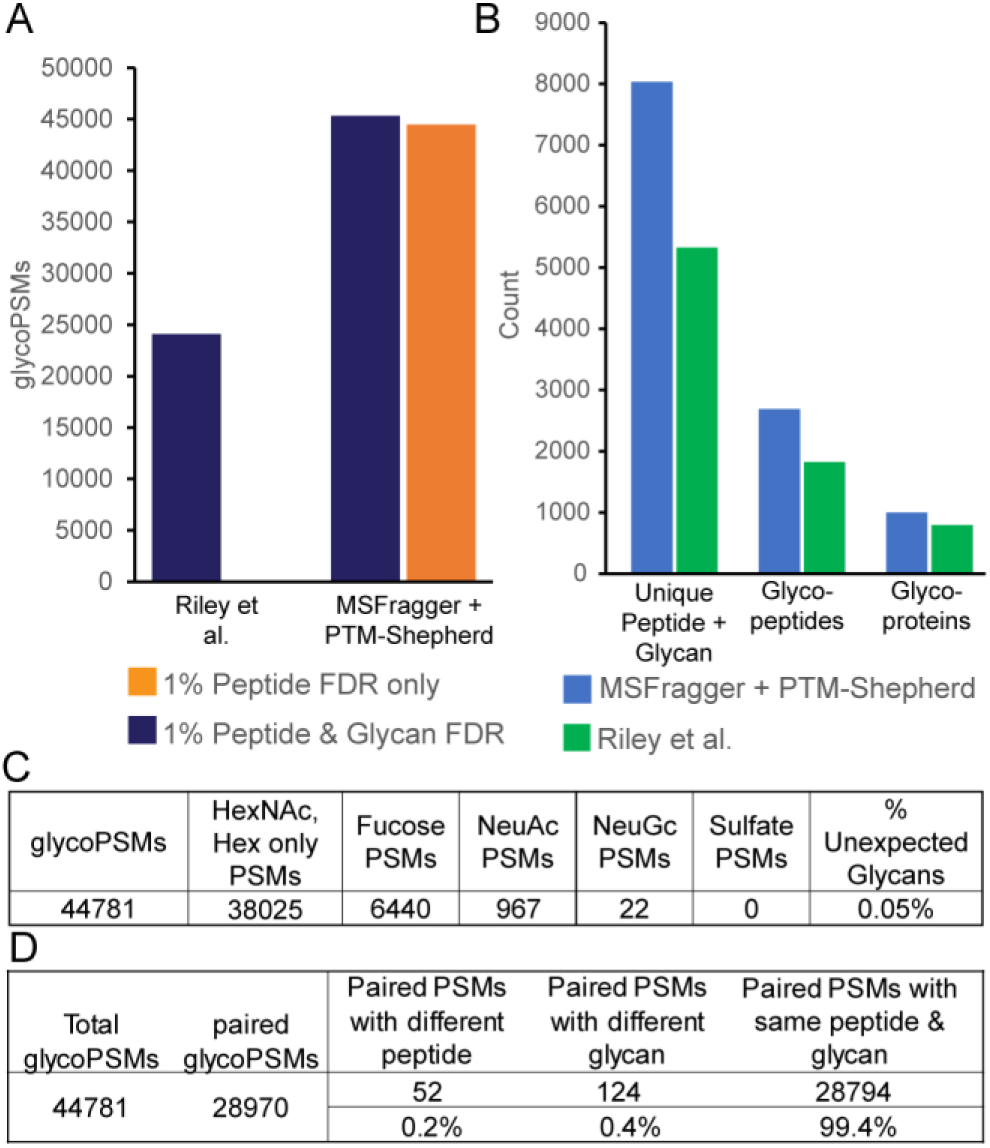
Comparison of glycoPSMs annotated from mouse brain tissue from Riley *et al.* A) GlycoPSMs annotated by the original analysis in Byonic and our re-analysis with MSFragger and PTM-Shepherd. The improvement in annotated spectra from MSFragger search is maintained even when applying 1% glycan FDR filtering. B) Unique glycoproteins, glycopeptide sequences, and glycan-peptide sequence combinations reported by MSFragger/PTM-Shepherd or Riley *et al*. C) Glycan composition categories observed sorted by residue type(s) contained, showing few unexpected (NeuGc or sulfate-containing) glycans. D) Analysis of paired HCD and AI-ETD scans of the same precursor from MSFragger/PTM-Shepherd search, assessed for whether the same peptide and glycan were matched in both of the paired scans.

Finally, while the entrapment glycan searches performed in yeast establish a strong foundation for FDR control in N-glycoproteomics analyses, the yeast glycoproteome contains few to no true glycan compositions very similar masses. To establish the performance of our method in more complex data, in which there are many examples of mass shifts that could be assigned to multiple possible glycan compositions expected to be present in the data, we re-analyzed N-glycopeptide data from five mouse tissue types originally presented in the description of the pGlyco2 software (9). In the MSFragger and PTM-Shepherd searches, we used the same glycan list as the “full list” from the yeast analysis, equivalent to the “pGlyco-N-mouse-large” glycan list from pGlyco3, allowing a single ammonium adduct. In all five tissue types, we find 40-66% more glycoPSMs at 1% peptide and glycan FDR than pGlyco3 when searching the same peptide and glycan lists, which in turn obtained a similar advantage over pGlyco2 (Fig. 5A). The pGlyco2 search (results as reported by (9)) did not include ammonium adducts as this feature was not supported in pGlyco2. As in the yeast data, we attribute the similarly sized increase in glycoPSMs annotated by MSFragger/PTM-Shepherd over pGlyco3 to the peptide-first search strategy and incorporation of additional information in glycan scoring as compared to pGlyco3. As in the other datasets, the increase in glycoPSMs translated to an increase in unique glycopeptide sequences in all tissue types and of glycoproteins in all tissues except liver and lung, in which glycoprotein counts were similar (Fig. 5B). We observed broadly similar trends in glycan compositions across tissue types, including the lack of NeuGc in brain tissue and predominance of high mannose glycans in liver (Fig. 5C). However, when we directly compare the frequency at which compositions containing various glycan residues were assigned, a clear pattern emerges of much higher rates of fucose-containing glycans assigned by pGlyco3 (orange bars) with roughly opposite increases in sialic acid-containing glycans assigned by PTM-Shepherd (green and yellow bars) (Fig. 5D). Given the high frequency of entrapment matches to fucosylated glycans by pGlyco3 in yeast, we suspect a similar bias towards fucose may be occurring in this mouse data and many of the fucosylated glycans assigned by pGlyco3 may in fact be sialylated, particularly since the substitution of two Fucoses for one NeuAc is a common glycan assignment mistake (30). An example spectrum with such a substitution is shown in Supplementary Figure S1, in which a series of abundant NeuAc-containing oxonium ions provides strong evidence for our composition assignment (HexNAc-4_Hex-5_NeuAc-1) over pGlyco3’s (HexNAc-4_Hex-5_Fuc-2). However, without a ground truth of known glycopeptides to compare against, we cannot say for certain which tool’s assignments are more frequently correct.

**Figure 5.**
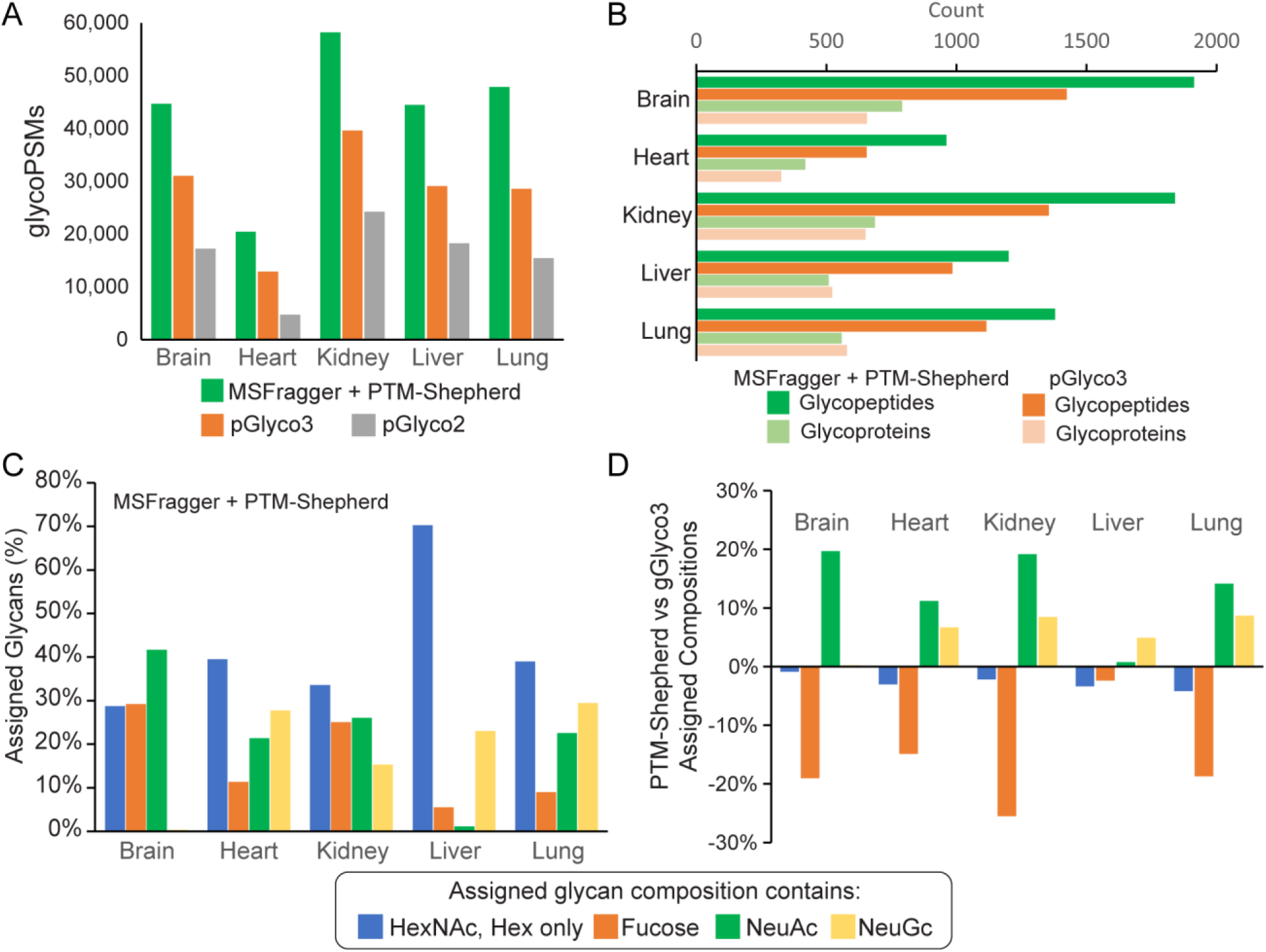
Comparison of N-glycan search in mouse tissue dataset between MSFragger + PTM-Shepherd and pGlyco3. A) GlycoPSMs passing 1% peptide and glycan FDR from MSFragger/PTM-Shepherd, pGlyco3, or pGlyco2 searches in each of 5 tissue types. B) Comparison of the number of unique glycoproteins and glycopeptide sequences (1% peptide and glycan FDR) from searches presented in A. C) PTM-Shepherd assigned compositions by residues included in the glycan. Note that glycoPSMs containing multiple residue types (e.g., Fucose and NeuAc) will be counted in multiple categories. D) Comparison of PTM-Shepherd assigned glycan compositions with pGlyco3 compositions. Positive values indicate more of a residue type assigned by PTM-Shepherd and negative values more assigned by pGlyco3. In all tissues, PTM-Shepherd assigns more sialic acid compositions while pGlyco3 assigned more Fucose.

Evaluating the accuracy of our method in this data is more challenging than in yeast, as there is not as straightforward a filter for entrapment compositions since glycans containing fucose, NeuAc, and NeuGc are all expected to be present in various tissues. However, we were able to generate a list of 248 entrapment glycans that have very similar masses (within 0.05 Da) of true mouse glycans using several substitutions of glycan residues (see Supplementary Data 1, “mouse entrapment” for the list of compositions). The majority of these entrapment glycans (180 compositions) contain phosphate, which is not expected to be present on glycans in these tissues, allowing entrapment compositions to be generated with similar masses to a wide range of mouse compositions. We also generated NeuAc (27), NeuGc (17), and Fuc (24)-containing entrapment glycans with similar masses to many of the most commonly observed mouse glycans. Unlike phosphate-containing entrapment glycans, these had to be manually checked against the mouse glycan list to ensure that true mouse glycans were not included as potential entrapments, resulting in much smaller numbers of these entrapment glycans compared to phosphate-containing ones. Nevertheless, by targeting these entrapment glycans to have similar masses to, and contain the same types of residues as, the most commonly observed real glycans we provide an intensive test of whether our method can distinguish between real and entrapment glycans in a complex analysis with many overlapping masses.

The results of entrapment glycan searches in each mouse tissue type are shown in Table 4. In each tissue, the rate of entrapment glycoPSMs matched remains at or below 1%, indicating good FDR control in a much more strenuous test than in the yeast data. Notably, almost no entrapment matches are made to phospho-glycans despite the presence of far more potential entrapment phospho-glycans than the other categories. The majority of entrapment glycoPSMs are to NeuAc in brain and kidney tissues and NeuGc in heart, lung, and liver, in each case corresponding to the most common category of glycan observed in the respective tissue. These observations, together with the lack of similar entrapment matches in the yeast data, suggest that the primary reason for the matches to entrapment glycans may be co-fragmentation of glycopeptides that results in oxonium ions from a glycan other than that of the precursor glycopeptide being including in scoring a given glycoPSM, increasing the rate of erroneous glycan assignments. This hypothesis is strongly supported by the brain tissue data, in which over 90% of all scans contain NeuAc oxonium ion(s) (despite only 40% of glycoPSMs being matched to NeuAc-containing compositions) while only ∼2% of scans have detectable NeuGc oxonium ion(s). Matches to NeuAc entrapment compositions were the vast majority of entrapment matches observed (89%) and no matches were made to NeuGc entrapment compositions (Table 4), despite being relatively common in the other tissue types where NeuGc oxonium ions occur with some frequency. Distinguishing the correct glycan from spectra in which oxonium ions seem to indicate an alternative composition is indeed a challenging problem, even for expert manual curation of glycan assignments. Maintaining a 1% entrapment glycan rate despite this extensive co-fragmentation is thus an impressive achievement for our method. Care should still be exercised in complex analyses, however, particularly when it is clear that oxonium ions for certain compositions are present in a majority of spectra. Narrower isolation windows, ion mobility filtering, and other measures to reduce co-fragmentation of glycopeptides would be likely to greatly reduce the incidence of this issue. It is reassuring to note that in all cases without such widespread oxonium presence, entrapment matches were nearly nonexistent.

**Table 4.** Results of entrapment glycan search in the mouse dataset. PSMs matched to entrapment glycans are shown as a count and percentage, remaining at or below the FDR used in the searches (1%). At right, entrapment glycoPSMs are sorted by glycan type. Details of the entrapment glycans used can be found in the supporting information.

| Sample | Total glycoPSMs | Entrapment glycoPSMs | Entrapment glycoPSMs (%) | Entrapment glycoPSMs by type: |  |  |  |
| --- | --- | --- | --- | --- | --- | --- | --- |
|  |  |  |  | NeuAc | NeuGc | Phospho | Fucose-only |
| Brain | 44931 | 491 | 1.1% | 436 | 0 | 0 | 55 |
| Heart | 20410 | 128 | 0.6% | 43 | 72 | 0 | 13 |
| Kidney | 65412 | 586 | 0.9% | 358 | 92 | 0 | 136 |
| Liver | 44476 | 24 | 0.1% | 3 | 13 | 2 | 6 |
| Lung | 47292 | 183 | 0.4% | 55 | 114 | 0 | 14 |

## Conclusions

We have developed a sensitive and robust method for determining the composition of the glycan component of N-glycopeptides from tandem MS data by combining information from multiple types of fragment ions with mass and isotope errors to distinguish between candidate compositions. We demonstrate that FDR control of the resulting glycan matches performs as expected even in analyses of complex glycan lists and in the presence of entrapment peptide sequences and glycan compositions. As glycoproteomics moves to larger and more complex samples, confident assignment of glycan compositions is critical to move beyond manual validation of glycopeptide spectra. We believe this work represents a promising step in this direction, enabling automated analysis of complex N-glycopeptide samples. Determining FDR for glycan assignment is a challenging problem with many complexities, and there remain several limitations in the method presented here, particularly in regard to searching O-glycopeptides, that we aim to address in future work. The probability ratios used to determine the likelihood of one glycan composition relative to another given the presence (or absence) of a particular fragment ion were determined empirically and are optimized for N-glycan fragmentation by HCD or stepped-energy HCD. While we have confirmed that these parameters still provide valid results for AI-ETD activation of N-glycans, probability parameters will likely need to be optimized specifically for other fragmentation methods to provide similarly high-quality results. Currently, all fragment ions of a given type are given the same probability ratios, despite the actual probabilities to observe various fragments varying greatly, and probabilities of observing the same fragment also depending on the precursor glycan. Using glycan and fragment-specific probabilities would likely improve the performance of glycan assignment, particularly when considering a wider range of fragmentation methods and settings. Recent reports have indicated that structure-specific fragmentation patterns could be used to infer glycan structure in addition to composition (33), a potential further application of fragment-specific probabilities. Finally, while the separation of glycan assignment from peptide sequencing greatly simplifies the problem and enables the high performance of our method, cases in which multiple glycans are present on a single peptide, or a combination of a glycan and other modification(s) found from open searching, are not currently supported and will require development of methods to localize multiple modification sites from these searches prior to glycan assignment. Methods for localizing multiple glycans on a peptide will also be required for the analysis of most O-glycopeptide data, as many O-glycopeptides contain multiple potential glycosites. While the method provided here is readily applicable to O-glycosylation in theory, handling multiply glycosylated peptides and tuning the fragment probabilities for O-glycans is needed to enable accurate O-glycan assignments.

## Acknowledgements

This work was funded in part by NIH grants R01-GM-094231 and U24-CA210967.

## Abbreviations

PTM: post-translational modification
NeuAc: N-acetyl neuraminic acid
NeuGc: N-glycolyl neuraminic acid
FDR: false discovery rate
PSM: peptide-spectrum match
glycoPSMs: glycopeptide-spectrum match
HCD: higher-energy C-trap dissociation
AI-ETD: activation ion-electron transfer dissociation
MS: mass spectrometry

## Data Availability

All raw data used can be found in the public repositories noted in the methods section. Processed results (PSM tables) used in the creation of all figures can be found at https://doi.org/10.5281/zenodo.5567044. PTM-Shepherd source code and a standalone JAR executable can be found at https://github.com/Nesvilab/PTM-Shepherd, and it is integrated into the FragPipe graphical user interface (http://fragpipe.nesvilab.org/).

## Author Contributions

D.A.P.: Conceptualization, Investigation, Software, Writing – Original Draft, Writing – Review & Editing. D.J.G.: Software, Methodology, Writing – Review & Editing. F. Y.: Software, Methodology, Writing – Review & Editing. A.I.N.: Conceptualization, Writing – Review & Editing, Supervision, Funding Acquisition

## Supplemental Data

This article contains supplemental data.

## References

1. Varki, A. (2017) Biological roles of glycans. Glycobiology 27, 3–49

2. Marsico, G., Russo, L., Quondamatteo, F., and Pandit, A. (2018) Glycosylation and Integrin Regulation in Cancer. Trends in Cancer 4, 537–552

3. Schedin-Weiss, S., Winblad, B., and Tjernberg, L. O. (2014) The role of protein glycosylation in Alzheimer disease. FEBS Journal 281, 46–62

4. York, I. A., Stevens, J., and Alymova, I. V. (2019) Influenza virus N-linked glycosylation and innate immunity. Portland Press Ltd

5. Thaysen-Andersen, M., Packer, N. H., and Schulz, B. L. (2016) Maturing glycoproteomics technologies provide unique structural insights into the N-glycoproteome and its regulation in health and disease. Molecular and Cellular Proteomics 15, 1773–1790

6. Suttapitugsakul, S., Sun, F., and Wu, R. (2019) Recent Advances in Glycoproteomic Analysis by Mass Spectrometry. Analytical Chemistry, acs.analchem.9b04651–acs.analchem.04659b04651

7. Reiding, K. R., Bondt, A., Franc, V., and Heck, A. J. R. (2018) The benefits of hybrid fragmentation methods for glycoproteomics. pp. 260–268, Elsevier B.V.

8. Cao, W., Liu, M., Kong, S., Wu, M., Zhang, Y., and Yang, P. (2021) Recent advances in software tools for more generic and precise intact glycopeptide analysis. Molecular & Cellular Proteomics 20, 100060–100060

9. Liu, M.-Q., Zeng, W.-F., Fang, P., Cao, W.-Q., Liu, C., Yan, G.-Q., Zhang, Y., Peng, C., Wu, J.-Q., Zhang, X.-J., Tu, H.-J., Chi, H., Sun, R.-X., Cao, Y., Dong, M.-Q., Jiang, B.-Y., Huang, J.-M., Shen, H.-L., Wong, C. C. L., He, S.-M., and Yang, P.-Y. (2017) pGlyco 2.0 enables precision N-glycoproteomics with comprehensive quality control and one-step mass spectrometry for intact glycopeptide identification. Nature Communications 8, 438–438

10. Zeng, W. F., Liu, M. Q., Zhang, Y., Wu, J. Q., Fang, P., Peng, C., Nie, A., Yan, G., Cao, W., Liu, C., Chi, H., Sun, R. X., Wong, C. C. L., He, S. M., and Yang, P. (2016) pGlyco: A pipeline for the identification of intact N-glycopeptides by using HCD- and CID-MS/MS and MS3. Scientific Reports 6

11. Zeng, W.-F., Cao, W.-Q., Liu, M.-Q., He, S.-M., Yang, and Peng, Y. (2021) Precise, Fast and Comprehensive Analysis of Intact Glycopeptides and Monosaccharide-Modifications with pGlyco3. bioRxiv

12. Bern, M., Kil, Y. J., and Becker, C. (2012) Byonic: Advanced Peptide and Protein Identification Software. Current Protocols in Bioinformatics 40, 13.20.11–13.20.14

13. Xiao, K., and Tian, Z. (2019) GPSeeker Enables Quantitative Structural N-Glycoproteomics for Site- And Structure-Specific Characterization of Differentially Expressed N-Glycosylation in Hepatocellular Carcinoma. Journal of Proteome Research 18, 2885–2895

14. Lu, L., Riley, N. M., Shortreed, M. R., Bertozzi, C. R., and Smith, L. M. (2020) O-Pair Search with MetaMorpheus for O-glycopeptide Characterization. bioRxiv, 2020.2005.2018.102327–102020.102305.102318.102327

15. He, L., Xin, L., Shan, B., Lajoie, G. A., and Ma, B. (2014) GlycoMaster DB: Software to assist the automated identification of N-linked glycopeptides by tandem mass spectrometry. Journal of Proteome Research 13, 3881–3895

16. Lynn, K.-S., Chen, C.-C., Lih, T. M., Cheng, C.-W., Su, W.-C., Chang, C.-H., Cheng, C.-Y., Hsu, W.-L., Chen, Y.-J., and Sung, T.-Y. (2015) MAGIC: An Automated N-Linked Glycoprotein Identification Tool Using a Y1-Ion Pattern Matching Algorithm and in Silico MS 2 Approach. Analytical Chemistry 87, 2466–2473

17. Polasky, D. A., Yu, F., Teo, G. C., and Nesvizhskii, A. I. (2020) Fast and comprehensive N- and O-glycoproteomics analysis with MSFragger-Glyco. Nature Methods 17, 1125–1132

18. Hu, Y., Shah, P., Clark, D. J., Ao, M., and Zhang, H. (2018) Reanalysis of Global Proteomic and Phosphoproteomic Data Identified a Large Number of Glycopeptides. Analytical Chemistry 90, 8065–8071

19. Hu, H., Khatri, K., Klein, J., Leymarie, N., and Zaia, J. (2016) A review of methods for interpretation of glycopeptide tandem mass spectral data. Glycoconjugate Journal 33, 285–296

20. Kawahara, R., Alagesan, K., Bern, M., Cao, W., Chalkley, R. J., Cheng, K., Choo, M. S., Edwards, N., Goldman, R., Hoffmann, M., Hu, Y., Huang, Y., Kim, J. Y., Kletter, D., Liquet-Weiland, B., Liu, M., Mechref, Y., Meng, B., Neelamegham, S., Nguyen-Khuong, T., Nilsson, J., Pap, A., Park, G. W., Parker, B. L., Pegg, C. L., Penninger, J. M., Phung, T. K., Pioch, M., Rapp, E., Sakalli, E., Sanda, M., Schulz, B. L., Scott, N. E., Sofronov, G., Stadlmann, J., Vakhrushev, S. Y., Woo, C. M., Wu, H.-Y., Yang, P., Ying, W., Zhang, H., Zhang, Y., Zhao, J., Zaia, J., Haslam, S. M., Palmisano, G., Yoo, J. S., Larson, G., Khoo, K.-H., Medzihradszky, K. F., Kolarich, D., Packer, N. H., and Thaysen-Andersen, M. (2021) Community Evaluation of Glycoproteomics Informatics Solutions Reveals High-Performance Search Strategies of Glycopeptide Data. bioRxiv

21. Keller, A., Nesvizhskii, A. I., Kolker, E., and Aebersold, R. (2002) Empirical statistical model to estimate the accuracy of peptide identifications made by MS/MS and database search. Analytical Chemistry 74, 5383–5392

22. Bollineni, R. C., Koehler, C. J., Gislefoss, R. E., Anonsen, J. H., and Thiede, B. (2018) Large-scale intact glycopeptide identification by Mascot database search. Scientific Reports 8

23. Fang, P., Xie, J. J., Sang, S., Zhang, L., Liu, M., Yang, L., Xu, Y., Yan, G., Yao, J., Gao, X., Qian, W., Wang, Z., Zhang, Y., Yang, P., and Shen, H. (2020) Multilayered N-Glycoproteome Profiling Reveals Highly Heterogeneous and Dysregulated Protein N-Glycosylation Related to Alzheimer’s Disease. Analytical chemistry 92, 867–874

24. Blazev, R., Ashwood, C., Abrahams, J. L., Chung, L. H., Francis, D., Yang, P., Watt, K. I., Qian, H., Quaife-Ryan, G. A., Hudson, J. E., Gregorevic, P., Thaysen-Andersen, M., and Parker, B. L. (2021) Integrated glycoproteomics identifies a role of N-glycosylation and galectin-1 on myogenesis and muscle development. Molecular and Cellular Proteomics 20, 100030–100030

25. Chen, Z., Yu, Q., Yu, Q., Johnson, J., Shipman, R., Zhong, X., Huang, J., Asthana, S., Carlsson, C., Okonkwo, O., and Li, L. (2021) In-depth site-specific analysis of N-glycoproteome in human cerebrospinal fluid (CSF) and glycosylation landscape changes in Alzheimer’s disease (AD). Molecular & cellular proteomics : MCP 0, 100081–100081

26. Hu, H., Khatri, K., and Zaia, J. (2017) Algorithms and design strategies towards automated glycoproteomics analysis. pp. 475–498

27. Hackett, W. E., and Zaia, J. (2021) The Need for Community Standards to Enable Accurate Comparison of Glycoproteomics Algorithm Performance. Molecules 26, 4757–4757

28. Hackett, W., and Zaia, J. (2021) Calculating glycoprotein similarities from mass spectrometric data. Molecular & Cellular Proteomics 20, 100028–100028

29. Darula, Z., and Medzihradszky, K. F. (2015) Carbamidomethylation Side Reactions May Lead to Glycan Misassignments in Glycopeptide Analysis. Analytical Chemistry 87, 6297–6302

30. Lee, L. Y., Moh, E. S. X., Parker, B. L., Bern, M., Packer, N. H., and Thaysen-Andersen, M. (2016) Toward Automated N-Glycopeptide Identification in Glycoproteomics. Journal of Proteome Research 15, 3904–3915

31. Zhu, Z., Su, X., Go, E. P., and Desaire, H. (2014) New glycoproteomics software, glycopep evaluator, generates decoy glycopeptides de novo and enables accurate false discovery rate analysis for small data sets. Analytical Chemistry 86, 9212–9219

32. Klein, J., Carvalho, L., Zaia, J., and Yy, J. (2021) Expanding N-Glycopeptide Identiications by Fragmentation Prediction and Glycome Network Smoothing.

33. Shen, J., Jia, L., Dang, L., Su, Y., Zhang, J., Xu, Y., Zhu, B., Chen, Z., Wu, J., Lan, R., Hao, Z., Ma, C., Zhao, T., Gao, N., Bai, J., Zhi, Y., Li, J., Zhang, J., and Sun, S. (2021) StrucGP: de novo structural sequencing of site-specific N-glycan on glycoproteins using a modularization strategy. Nature Methods

34. Yu, F., Teo, G. C., Kong, A. T., Haynes, S. E., Avtonomov, D. M., Geiszler, D. J., and Nesvizhskii, A. I. (2020) Identification of modified peptides using localization-aware open search. Nature Communications 11, 4065–4065

35. Geiszler, D. J., Kong, A. T., Avtonomov, D. M., Yu, F., da Veiga Leprevost, F., and Nesvizhskii, A. I. (2021) PTM-shepherd: Analysis and summarization of post-translational and chemical modifications from open search results. Molecular and Cellular Proteomics 20, 100018–100018

36. Deutsch, E. W., Csordas, A., Sun, Z., Jarnuczak, A., Perez-Riverol, Y., Ternent, T., Campbell, D. S., Bernal-Llinares, M., Okuda, S., Kawano, S., Moritz, R. L., Carver, J. J., Wang, M., Ishihama, Y., Bandeira, N., Hermjakob, H., and Vizcaíno, J. A. (2017) The ProteomeXchange consortium in 2017: Supporting the cultural change in proteomics public data deposition. Nucleic Acids Research 45, D1100–D1106

37. Kessner, D., Chambers, M., Burke, R., Agus, D., and Mallick, P. (2008) ProteoWizard: Open source software for rapid proteomics tools development. Bioinformatics 24, 2534–2536

38. Riley, N. M., Hebert, A. S., Westphall, M. S., and Coon, J. J. (2019) Capturing site-specific heterogeneity with large-scale N-glycoproteome analysis. Nature Communications 10, 1311–1311

39. da Veiga Leprevost, F., Haynes, S. E., Avtonomov, D. M., Chang, H. Y., Shanmugam, A. K., Mellacheruvu, D., Kong, A. T., and Nesvizhskii, A. I. (2020) Philosopher: a versatile toolkit for shotgun proteomics data analysis. Nature Methods 17, 869–870

40. Nesvizhskii, A. I., Keller, A., Kolker, E., and Aebersold, R. (2003) A statistical model for identifying proteins by tandem mass spectrometry. Analytical Chemistry 75, 4646–4658

41. Medzihradszky, K. F., Kaasik, K., and Chalkley, R. J. (2015) Characterizing sialic acid variants at the glycopeptide level. Analytical Chemistry 87, 3064–3071

42. Halim, A., Westerlind, U., Pett, C., Schorlemer, M., Rüetschi, U., Brinkmalm, G., Sihlbom, C., Lengqvist, J., Larson, G., and Nilsson, J. (2014) Assignment of saccharide identities through analysis of oxonium ion fragmentation profiles in LC-MS/MS of glycopeptides. Journal of Proteome Research 13, 6024–6032

43. Pett, C., Nasir, W., Sihlbom, C., Olsson, B. M., Caixeta, V., Schorlemer, M., Zahedi, R. P., Larson, G., Nilsson, J., and Westerlind, U. (2018) Effective Assignment of α2,3/α2,6-Sialic Acid Isomers by LC-MS/MS-Based Glycoproteomics. Angewandte Chemie - International Edition 57, 9320–9324

44. Ács, A., Ozohanics, O., Vékey, K., Drahos, L., and Turiák, L. (2018) Distinguishing Core and Antenna Fucosylated Glycopeptides Based on Low-Energy Tandem Mass Spectra. Analytical Chemistry 90, 12776–12782

45. Lakbub, J. C., Su, X., Hua, D., Go, E. P., and Desaire, H. (2018) Dissecting the dissociation patterns of fucosylated glycopeptides undergoing CID: A case study in improving automated glycopeptide analysis scoring algorithms. Analytical Methods 10, 256–262

46. Caval, T., Zhu, J., Tian, W., Remmelzwaal, S., Yang, Z., Clausen, H., and Heck, A. J. R. (2019) Targeted analysis of lysosomal directed proteins and their sites of mannose-6-phosphate modification. Molecular and Cellular Proteomics 18, 16–27

47. Kuo, C. W., Guu, S. Y., and Khoo, K. H. (2018) Distinctive and Complementary MS 2 Fragmentation Characteristics for Identification of Sulfated Sialylated N-Glycopeptides by nanoLC-MS/MS Workflow. Journal of the American Society for Mass Spectrometry 29, 1166–1178

48. Sanda, M., Benicky, J., and Goldman, R. (2020) Low Collision Energy Fragmentation in Structure-Specific Glycoproteomics Analysis. Analytical Chemistry 92, 8262–8267

49. Yu, J., Schorlemer, M., Gomez Toledo, A., Pett, C., Sihlbom, C., Larson, G., Westerlind, U., and Nilsson, J. (2016) Distinctive MS/MS Fragmentation Pathways of Glycopeptide-Generated Oxonium Ions Provide Evidence of the Glycan Structure. Chemistry - A European Journal 22, 1114–1124

50. Hoffmann, M., Pioch, M., Pralow, A., Hennig, R., Kottler, R., Reichl, U., and Rapp, E. (2018) The Fine Art of Destruction: A Guide to In-Depth Glycoproteomic Analyses—Exploiting the Diagnostic Potential of Fragment Ions. Proteomics 18

51. Yang, Y., Cao, W., Yan, G., Kong, S., Wu, M., Yang, P., and Qiao, L. (2021) GproDIA enables data-independent acquisition glycoproteomics with comprehensive statistical control. bioRxiv, 2021.2003.2020.436117–432021.436103.436120.436117

52. Trinidad, J. C., Schoepfer, R., Burlingame, A. L., and Medzihradszky, K. F. (2013) N- and O-Glycosylation in the murine synaptosome. Molecular and Cellular Proteomics 12, 3474–3488

