## Supplementary Information for "Multi-attribute Glycan Identification and FDR Control for Glycoproteomics"

Table of Contents:

1. Table S1: Probability ratios for Y and oxonium ion types and peakpicking errors
2. Table S2: Peakpicking error probabilities used to generate peakpicking error score
3. Table S3. Glycan lists used in MSFragger searches and PTM-Shepherd glycan assignments for all datasets
4. Table S4. Glycans assigned naively by closest match to input glycan list in search of yeast dataset
5. Table S5. GlycoPSMs with entrapment (mouse) peptides in searches of yeast data
6. Table S6. Differences in assigned glycans in the Riley et al. dataset
7. Figure S1. Example spectrum from mouse data comparing NeuAc vs Fuc-2 assignment
8. Supplementary data 1: lists of glycan compositions used in all searches (.xlsx)

Table S1. Probability ratios for various fragment ion types. For each fragment ion type, probability ratios used for all analyses reported here are displayed. Scores are computed as the summed log of these ratios for each fragment ion considered, thus, a probability ratio of 1 has no effect on the score, ratios greater than 1 improve the score, and ratios less than 1 reduce the score. For oxonium ions, an expected intensity (relative to the spectrum base peak) is provided to reduce the impact of ions resulting from co-fragmentation of another glycopeptide. The hit ratio is multiplied by the ratio of observed / expected intensity. If observed intensity is much lower than expected intensity, such that it drives the resulting product to be less than 1, the score is set to 1 for that fragment to prevent such cases from actively reducing the score (such cases will have no effect on score). Note that fucose-containing oxonium ions have reduced penalty for misses and lower expected intensity, as core-fucosylated glycans may not produce such ions.

| Ion Type | Residue Category | Found ("Hit") Ratio | Not Found ("Miss") Ratio | Expected Intensity (%) |
| --- | --- | --- | --- | --- |
| Y | HexNAc/Hex | 5 | 0.5 | n/a |
| Y | HexNAc/Hex with Fucose | 2 | 0.5 | n/a |
| Oxonium | NeuAc | 2 | 0.05 | 20% |
| Oxonium | NeuGc | 2 | 0.05 | 20% |
| Oxonium | Fucose | 2 | 0.5 | 10% |
| Oxonium | Phosphate | 2 | 0.05 | 20% |
| Oxonium | Sulfate | 2 | 0.05 | 20% |

Table S2. Probability values for peakpicking errors. The probability ratio for pairwise comparisons is the log of the ratio of the values below for the peakpicking error of each candidate. For example, if candidate 1 has no peakpicking error and candidate 2 has +2 peakpicking error, the peakpicking error score would be log( 1 / 0.5 ).

| Peakpicking Error | Probability value |
| --- | --- |
| -2 | 0.125 |
| -1 | 0.25 |
| 0 | 1 |
| 1 | 0.95 |
| 2 | 0.5 |
| 3 | 0.25 |
| 4 | 0.125 |

Table S3. Glycan lists used in MSFragger searches and PTM-Shepherd glycan assignments for all datasets. Glycan composition lists are enumerated in the supplementary data 1 sheet. The pGlyco-1670 list was searched with ammonium adducts in MSFragger, resulting in 2325 unique masses. The number after the dash corresponds to the number of unique compositions searched in the list.

| Name/Reference in manuscript | Associated Figure(s) | Associated Table(s) | Glycan (mass offset) List used in MSFragger Search | Glycan List used in PTM-Shepherd Glycan Assignment |
| --- | --- | --- | --- | --- |
| yeast data, mouse list | Fig. 2B, far left | Table 1 (top row) | mouse-182 | pGlyco-1670 |
| yeast data, yeast list | Fig. 2B, 2nd from left | Table 1 (2nd row) | yeast-186 | pGlyco-1670 |
| yeast data, full list | Fig. 2B, middle, Figure 3 | Table 1 (3rd row), Table 2, Table 3 | pGlyco-1670 | pGlyco-1670 |
| Riley | Figure 4 |  | mouse-182 | PTM-Shepherd internal plus O-glycans-2335 |
| mouse 5-tissue (base) | Figure 5 |  | pGlyco-1670 | pGlyco-1670 |
| mouse 5-tissue (entrapment) |  | Table 4 | pGlyco-1670 | mouse entrapment-2029 |

Table S4: Glycans assigned naively by closest match to input glycan list in search of yeast dataset. Entrapment peptides included in both searches, entrapment glycans for “mouse list” search but not for “yeast list” search. Asterisk (*) indicates no glycans were matched to a given composition type because no such compositions are present in the list being searched and matched against. When entrapment glycans are searched, naïve mass matching without FDR control matches a large number of entrapment glycans. Searches without entrapment glycans do match any (by definition), but the actual rate of false matching is unknown unless glycan assignment is FDR controlled.

| Search Engine | Glycan assignment | Glycan list searched | Reported glycoPSMs | NeuAc-containing PSMs | NeuGc-containing PSMs | Total % suspect |
| --- | --- | --- | --- | --- | --- | --- |
| MSFragger | (closest mass) | Mouse list (182) | 4015 | 873 | 0* | 22% |
| MSFragger | (closest mass) | Yeast list (186) | 8374 | 0* | 0* | 0%* |

Table S5: GlycoPSMs with entrapment (mouse) peptides in searches of yeast data. GlycoPSM counts for mouse and yeast peptides from searches of combined yeast/mouse database with various glycan lists (same data as in Figure 2), filtered to 1% peptide and 1% glycan FDR. Entrapment (mouse) peptide PSM rate remains 1% or less in all cases. When considering all PSMs (i.e. including non-glyco PSMs), the entrapment peptide match rate drops to ~0.1% for all MSFragger/PTM-Shepherd results. pGlyco 3 data shown in comparison taken from (1) and used same peptide and glycan databases in search.

| Search Engine | Glycans Searched | glycoPSMs w/ yeast peptide | glycoPSMs w/ mouse peptide | Total glycoPSMs | Mouse peptide % |
| --- | --- | --- | --- | --- | --- |
| MSFragger+PTM-S | Mouse list (182) | 3164 | 6 | 3170 | 0.2% |
| MSFragger+PTM-S | Yeast list (186) | 7572 | 26 | 7598 | 0.3% |
| MSFragger+PTM-S | Full Database (1670) | 8188 | 46 | 8234 | 0.6% |
| pGlyco3 | Full Database (1670) | 3398 | 7 | 3405 | 0.2% |

Table S6. GlycoPSMs from the Riley et al. data with different glycan assignments from MSFragger/PTM-Shepherd and Byonic. Several of the most common differentially assigned compositions are shown. By far the most common assignment difference is HexNAc(2)Hex(n) from PTM-Shepherd vs HexNAc(6)Hex(n-5) from Byonic. Asterisk (*) indicates that the Byonic assigned composition is 15.99 Da lighter than the PTM-Shepherd assigned composition, which is balanced by the addition of an oxidized Met on the peptide in the Byonic result.

| PTM-Shepherd Glycan | Riley et al. (Byonic) Glycan | PSM Count |
| --- | --- | --- |
| HexNAc(2)Hex(8) | HexNAc(6)Hex(3) | 179 |
| HexNAc(2)Hex(9) | HexNAc(6)Hex(4) | 92 |
| HexNAc(4)Hex(5)Fuc(1)NeuAc(2) | HexNAc(4)Hex(5)Fuc(3)NeuAc(1) | 13 |
| HexNAc(2)Hex(5) | HexNAc(2)Hex(4)Fuc(1)* | 7 |
| HexNAc(4)Hex(6)NeuAc(1) | HexNAc(4)Hex(6)Fuc(2) | 5 |

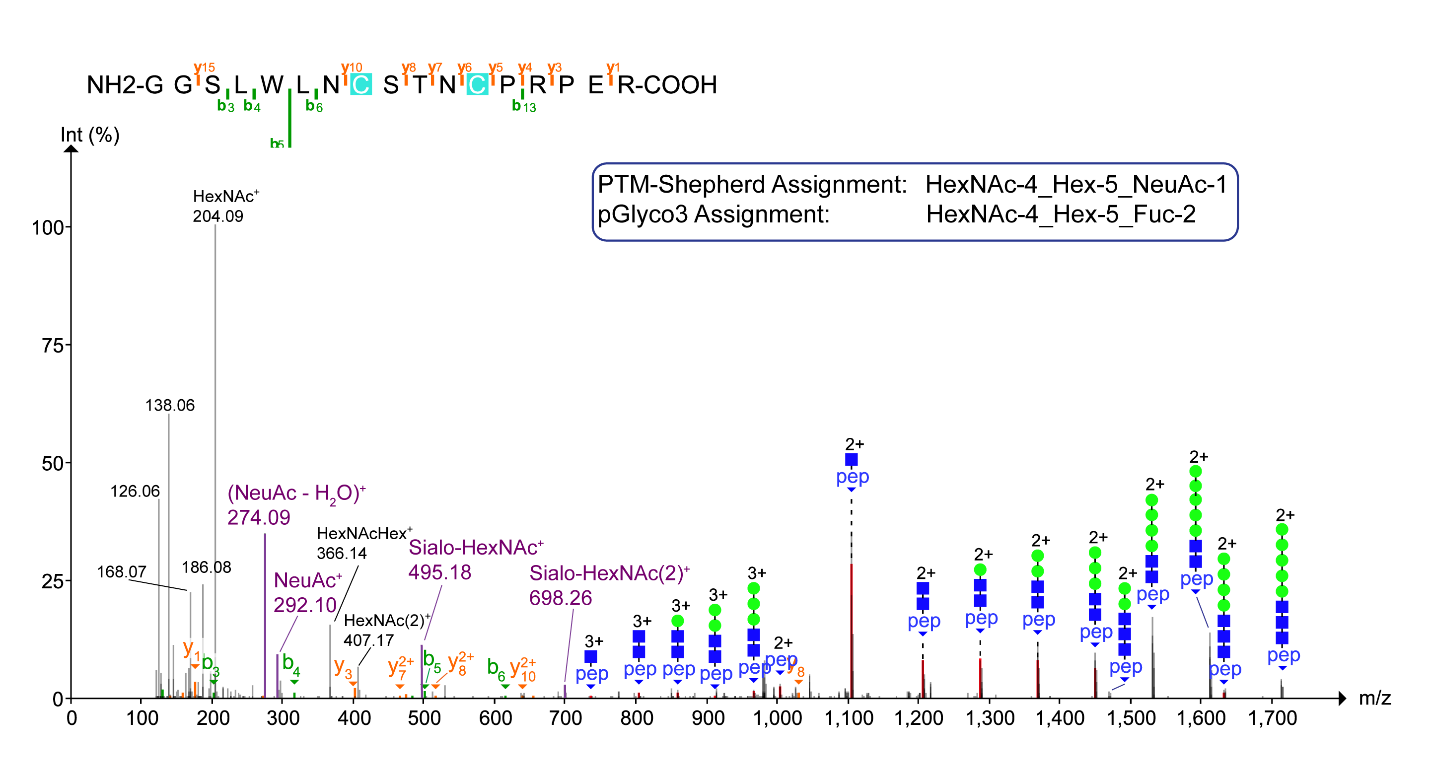

Figure S1. Example spectrum from mouse data comparing NeuAc vs Fuc-2 assignment. Our method and pGlyco assign the same peptide sequence but glycan compositions differing by NeuAc (our method) vs 2 Fucoses (pGlyco3). No Y or oxonium ions containing Fucose are observed in the spectrum, but an abundant series of NeuAc oxonium ions (purple) is observed.

References

1.	Zeng, W.-F., Cao, W.-Q., Liu, M.-Q., He, S.-M., Yang, and Peng, Y. (2021) Precise, Fast and Comprehensive Analysis of Intact Glycopeptides and Monosaccharide-Modifications with pGlyco3. *bioRxiv*
